# Socially transmitted knowledge of hibernation sites in bats

**DOI:** 10.64898/2026.08.06.743314

**Authors:** Simon Ripperger, Gerald G. Carter, Lutz Ittermann, Jörg Harder, Brigitte Kaltofen, Robert Henning, Karsten Dedek, Paul Voigt, Ahana Aurora Fernandez

**Author notes:** equal contributions (Simon Ripperger, Gerald G. Carter, Ahana Aurora Fernandez).

## Abstract

In temperate regions around the world, bats travel long distances every winter to gather at hibernation sites. A longstanding hypothesis is that each new generation of bats learns about the locations of these sites (called ‘hibernacula’) from older individuals, yet clear and compelling evidence demonstrating social transmission of this knowledge has been lacking. Here, we compiled 30,882 observations from 1985 to 2023 of 13,852 Greater mouse-eared bats (*Myotis myotis*) that were banded and observed at summer roosts, winter hibernacula, or both. Our analyses revealed four lines of evidence that Greater mouse-eared bats find suitable hibernacula using social information acquired at summer roosts. First, naïve yearlings were more likely to be seen sharing their first hibernacula with adults from their summer birth colony relative to a null model where bats moved independently. Second, adult bats were also more likely to co-switch together into the same hibernacula across winters than expected from independent movements. Third, bats that roosted together in the summer were more likely to share a different site as a hibernaculum during the winter: being observed together during a summer changed the probability of a pair being observed together during a winter from 5% to 12%. Finally, high-resolution tracking revealed an instance of tandem flights to hibernacula sites during the summer, demonstrating that yearlings can learn from experienced adult bats months before hibernation. Together, our findings show that maternity colonies serve as “information centers” where females acquire knowledge of suitable hibernation sites throughout their long lives.

## RESULTS AND DISCUSSION

### Four decades of bat banding show bats switch among multiple summer and winter roosts

In most temperate bat species, females communally rear their young in summer “maternity colonies” and both males and females move from summer sites to winter hibernacula, which can be dozens or even hundreds of kilometers away. A long-standing question is whether bats’ knowledge of hibernation sites is socially transmitted. To answer this question, we combined our own observations with those obtained from the center for bat banding at the Saxon State Office for Environment, Agriculture and Geology in Dresden, Germany, to compile 30,882 observations of 13,852 Greater mouse-eared bats (*Myotis myotis*) that were individually marked (with a metal band on their forearm) over a period of 38 years. Of these banded bats, 6412 were seen more than once, and 184 bats were seen 10 or more times (Fig. S1).

Bats were observed at roost sites (n=351) that were used as hibernacula in the winter, maternity (breeding) colonies or non-maternity colonies in the summer (Fig. S2), or as sites for “swarming”—a behavior (coinciding with mating in some bat species) where many bats aggregate at hibernacula during the fall prior to hibernation^1–5^. Most roost sites (n=264) served a single function, but others were used in more than one context. Bats switched between hibernation sites across winters, but many winter sites were not known or sampled. The average probability of a banded bat being observed in a winter was therefore only 36.8% (n=13,582). Each bat was observed at 1-5 of the sampled hibernation sites, and the number of sites used by a bat depended on the number of times it was observed (Fig. S2).

For all bats with known birth sites, the mean distances between birth sites and hibernation sites sampled over their lifetime was 33 km for females (bootstrapped 95% confidence interval=[31,36]) and 39 km for males ([37,42]). The maximum observed distances were 246 km for females and 235 km for males. Mapping these distances from summer birth sites to winter hibernation sites shows that bats were not limited in which sampled sites they could use; all sampled summer and winter sites were close enough to be used by any bat (Fig. 1).

**Figure 1.**
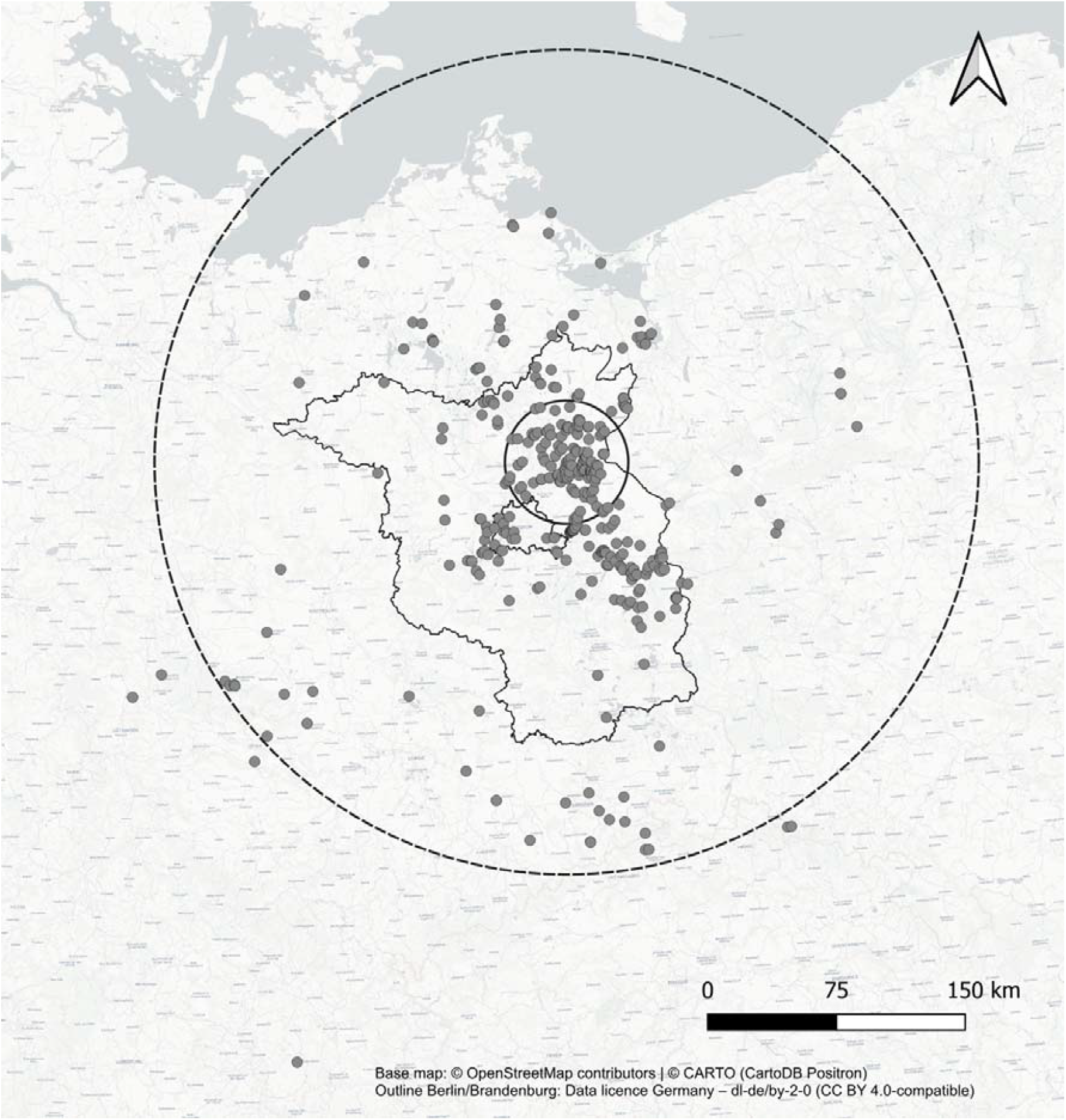
Locations of roost sites. Most roosting sites are located in the states of Berlin and Brandenburg, Germany (outlined in black). Circles centered around the maternity colony where bats displayed in Figure 3 were tagged for tracking show the mean observed distance (radius of small solid circle) and maximum observed distance (radius of large dashed circle) between birth sites and hibernacula.

Hibernacula are used by both sexes, but maternity colonies are almost exclusively females and their offspring (Fig. S3). If social information transfer occurs at maternity sites, we might expect a female bias in the frequency of discovery and switching of hibernation sites. Indeed, we found that females switched hibernacula more often than males (Fig. S4, average probability of observing a switch from a Bayesian hurdle Poisson model: all bats: 0.51, 95% Highest Density Interval (HDI)=[0.49, 0.53]; females=0.58 [0.56, 0.60], males=0.45 [0.43, 0.47]); see methods for details). Next, we examined what factors predict discovery and use of a bat’s first hibernaculum.

### The first hibernation sites used by female yearling bats are not the closest to their birth site

Yearling (young-of-the-year) bats are naïve to locations of hibernacula. One hypothesis for how yearlings find hibernation sites is that they sample or search the areas near their birth site, perhaps by detecting cues of swarming^1–5^. If so, then we might expect the first hibernation sites used by yearlings to be closer to their birth site relative to their second hibernation site because the search time and radius should increase over time. For completely random and continuous search, the probability of an earlier hibernacula being closer to its origin approaches 100% as search time goes on.

We found that first hibernation sites were closer to their birth site in only 72% of cases with male bats (95% HDI=56% to 87%, Bayesian Bernoulli model). In females, however, the probability of hibernating closer to their birth site during their first winter was surprisingly low— only 29% (95% HDI=[12, 50%]). Contrary to the prediction of finding hibernacula near birth sites, females were more likely to hibernate at sites that were *farther* from their birth site as yearlings than as adults. This finding suggests that bats, and especially females, are not discovering hibernacula through independent sampling of their environment.

### Naïve yearlings shared their first hibernacula with adults from their summer birth colony more often than expected from a null model of bats moving independently

An alternative hypothesis is that yearlings learn about suitable hibernation sites from adults at their summer maternity colony. In past studies, genetic identification of mother-pup pairs at swarming sites has been at least consistent with the hypothesis that some yearlings learned about swarming sites from their mothers^5^, but previous analyses of associations of tagged bats during swarming (n=614 bats^4^), and analyses of the age and sex ratios of captured adults and juveniles (e.g. Piksa ^3^ and studies cited herein) failed to find clear support for social information transfer.

In Greater mouse-eared bats, births occur during June and July, adult females leave maternity roosts end of July to early August, mating peaks between mid-August and mid-September, and bats move to hibernacula mid-September to mid-October, with adult females leaving the maternity colony soon after their offspring has fledged, causing summer colonies to be occupied only by juveniles^6–9^. However, there is ample opportunity for yearlings to learn locations of hibernation sites from adult roostmates for the months they are together before hibernation. If so, we expected to observe yearlings hibernating with at least one adult bat from their summer birth site, and we expected to observed these events more often than expected from a null model based on non-social independent movements.

We found the first direct evidence that yearling bats learn about hibernacula from adult members of their colonies. Of the 7826 yearlings with known summer birth sites, we observed 931 yearlings move from a summer site (maternity colony) to their first winter hibernation site. Of these 931 movements, we observed 364 cases (39%) of where a yearling and adult from the same summer site made the same movement (i.e. a co-switch). This rate of co-switching was greater than expected from a null model simulating independent movements without social information transfer (p<0.001, 95% of expected values from null model=[36.5, 37.8%], see Methods). We observed the same result when testing only female yearlings (observed=0.42, p<0.001, 95% of expected proportions between 0.39 and 0.41) and only male yearlings (observed=0.36, p<0.001, 95% of expected proportions between 0.34 and 0.35).

### Co-switching of hibernacula was more likely than expected by chance

We found that social learning is not limited to yearling bats. Co-switching between hibernation sites by adult bats across winters also occurred more often than expected from a null model. We tested this by first gathering the 1,040 winter observations of co-roosting pairs of banded bats that were observed at more than one hibernaculum, and then counting cases where the pair was seen in the same hibernation site on one date and then seen again together at a different hibernaculum on a later date. We found 34 cases, which was twice as often as expected from a null model where we randomized the observation dates of winter sites within each bat (p < 0.001, expected cases=17.6; 95% quantiles=[12, 24%]).

### Summer co-roosting predict co-roosting at winter hibernation sites

If adults learn about hibernation sites by following conspecifics during the summer, then an observation of two bats together in the summer should increase the probability of those bats sharing a hibernaculum during the winter. To test this hypothesis, we created weighted networks of summer and winter co-roosting rates using the 180 bats that were seen at least 3 times in the winter and at least 3 times in the summer. To estimate the extent to which winter co-roosting probabilities increased with observed summer co-roosting rates, we fit three Bayesian multi-membership models. All three models revealed that summer co-roosting predicted co-roosting at different winter sites (Fig. 2).

**Figure 2.**
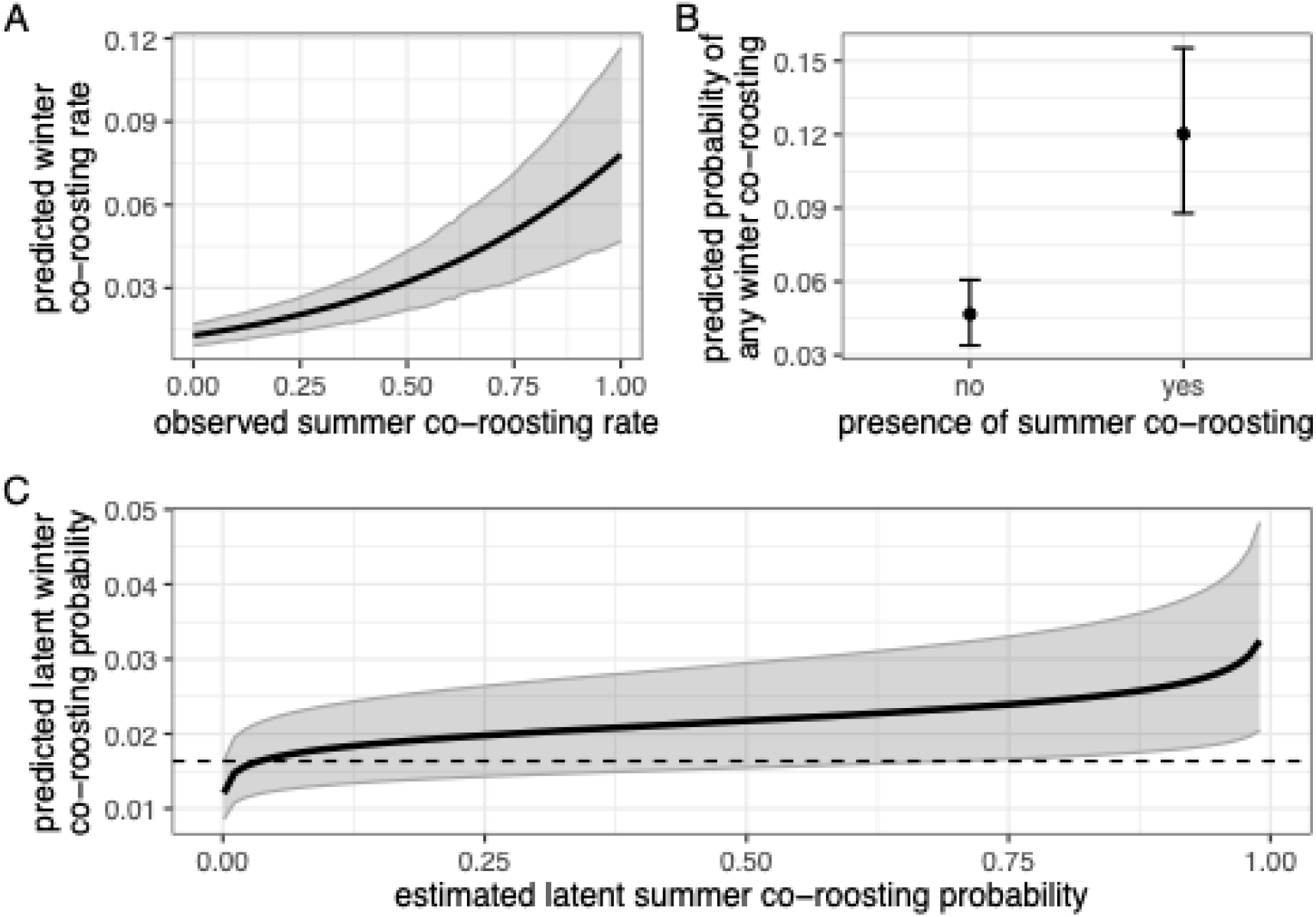
Summer co-roosting predicts winter co-roosting. Panels show results from three alternative Bayesian models. Panel A shows predicted probability of winter co-roosting based on observed summer co-roosting rates (proportions of actual and possible counts) from the first model. Panel B shows predicted probability of winter co-roosting based on presence or absence of summer co-roosting rates (after conditioning on sampling effort) from the second model. Panel C shows predicted latent probabilities of winter co-roosting by estimated latent probability of summer co-roosting from the third model. Shaded regions and error bars show 95% credible intervals. Posterior distributions are shown in Fig. S6–S8.

For the first beta-binomial model, the outcome was the count of winter co-roosting observations out of the total number of winter opportunities to co-roost, the predictor was the observed summer co-roosting rate (the proportion of summer opportunities in which the dyad was observed together), and a multi-membership group effect accounted for the non-independence of dyads sharing one or both bats. We used default uninformative priors. Rhat values for model parameters were 1.00-1.01 and effective sample sizes were 837 to 4061. According to the first model, the observed rate of summer co-roosting clearly predicted winter co-roosting (Fig. 2a, S5, Table S1), with a summer co-roosting rate of 0% predicting a winter co-roosting rate of 1% (95% credible interval (CrI)=[0.9, 1.7%]), and a summer co-roosting rate of 100% predicting a winter co-roosting rate of 7.8% (95% CrI=[4.7, 11.7%]).

For the second model, we asked whether pairs seen together during any sampled summer (true/false) were more likely to be seen together during any sampled winter (Bernoulli outcome). To control for sampling effort in this model, we included log counts of possible summer co-roosting and log counts of possible winter co-roosting for each dyad as covariates. We used default uninformative priors. Rhat values for model parameters were 1.00 and effective sample sizes were 3795 to 21145. According to the second model, pairs never observed together during the summer had a 4.7% chance (95% CrI=[3.4, 6.1%]) of ever being seen together during the winter, whereas a pair of bats seen together during the summer had a 12% chance (95% CrI=[8.8, 15.5%]) of being seen together at least once during the winter (Fig. 2b, S6).

To account for uncertainty in estimated summer co-roosting rates, we used the bisonR package^10^ to propagate uncertainty in summer co-roosting into a regression predicting winter co-roosting. For priors on summer co-roosting rates, we used a normally distributed prior centered at 3 on the logit scale (corresponding to an average co-roosting probability of 4.7%) with a standard deviation of 3, indicating that we expect most dyads to have low co-roosting probabilities but with very high uncertainty. We found that the predicted latent rates of summer and winter co-roosting were associated, with a latent summer co-roosting probability of 0.1% predicting a winter co-roosting probability of 1.2% (95% CrI=[0.9, 1.6%]) and a summer co-roosting probability of 99% predicting a winter co-roosting probability of 3.2% (95% CrI=[2.0, 4.8%]; Fig. 2c, S7).

### Social information transfer occurs through tandem flights

Social transmission of knowledge of winter hibernacula requires a mechanism of social information transfer, such as tandem flights^5,11–13^. Bats flying in tandem with their offspring over distances of several hundred meters provide direct evidence for information transfer from mothers to offspring for movements between summer roosts^14^. This “maternal guidance hypothesis” has long been discussed as a possible mechanism for young bats to find hibernation sites^3,5^. However, observing such co-movements is difficult.

Using high-resolution automated telemetry, we were fortunate to observe striking direct evidence for the maternal guidance hypothesis. In 2022, we fitted 40 bats with sensor nodes to track their presence at seven hibernacula equipped with tracking stations. We detected one tagged adult and two tagged juveniles from the same roost arriving at automated telemetry stations at different hibernacula within seconds or minutes during two different nights (Figure 3). One of the yearlings shared at least one allele at all loci that amplified in both individuals (15 out of 20 loci) and no mismatch was found. The second yearling shared alleles at three out of 12 loci. We, therefore, identify the three bats as mother, yearling offspring, and an unrelated yearling. During five occasions in late July and early August, the trio was detected at four of the seven hibernacula (Fig. 3, S8). Received signals of the arriving mother-pup pair were received only 4s, 15s, 26s, 39s, and 90s apart. The two bats stayed at a site for 6-72 minutes, before departing either simultaneously or within 33 s of each other (Fig. 3). Signals of the unrelated yearling were received along with signals of the mother-pup pair, but the timing of the visits was less synchronized (Fig. 3). These observations of the same pairs of bats repeatedly arriving at and departing from hibernacula within short time periods after resting for prolonged time (during two different nights and at four different locations) implicate tandem flights during the summer as a mechanism for naïve yearlings to gather information on the location of hibernacula. Yearlings can be introduced to several hibernacula during the first summer months of their lives.

**Figure 3.**
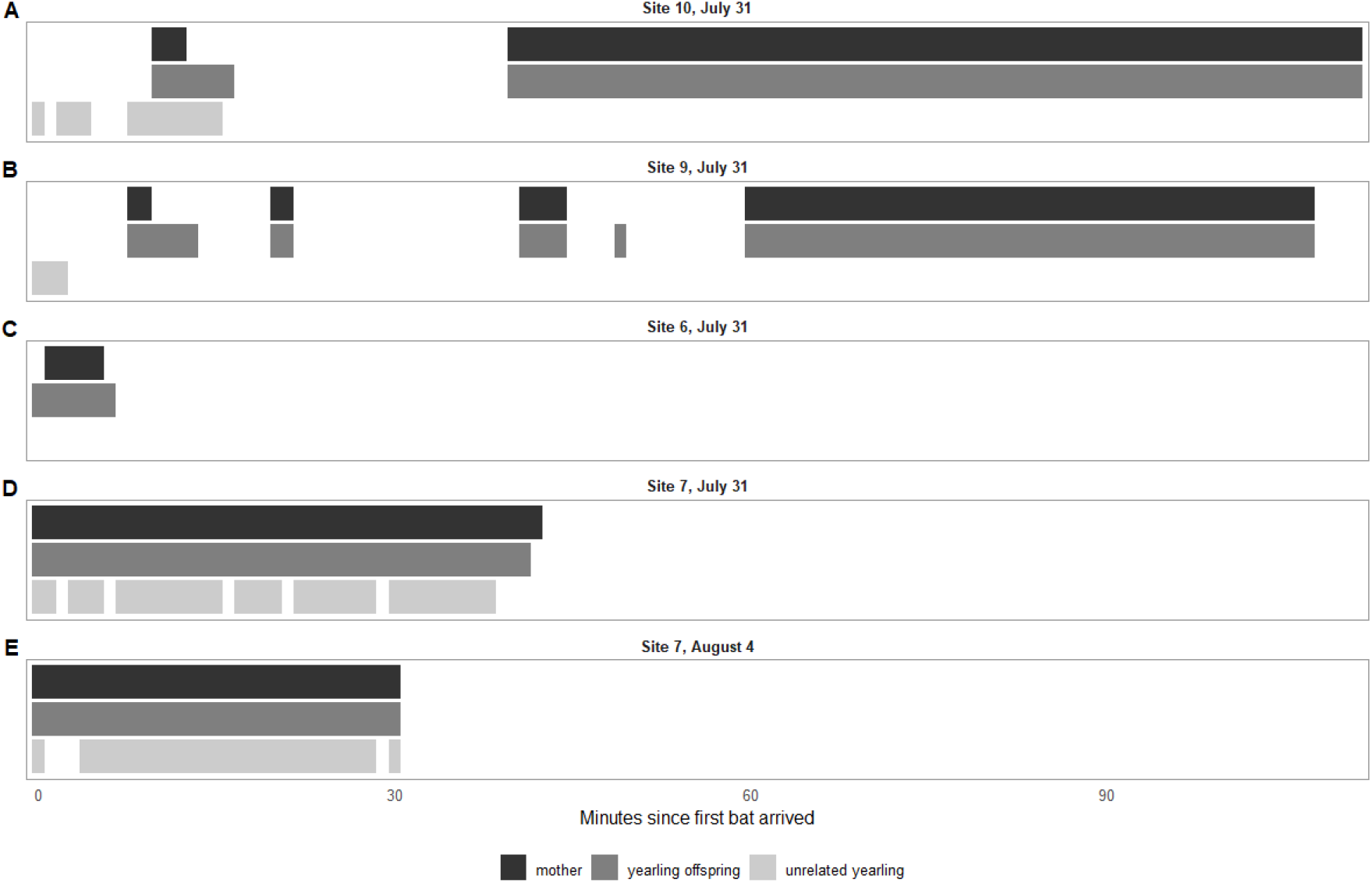
Timing of visits to hibernacula by three tagged bats during summer. Horizontal bars show the presence and co-occurrences of a mother (black), her yearling offspring (dark grey) and an unrelated yearling (light grey) at four sites on five days. Site numbers refer to Figure S9.

Taken together, our findings suggest that yearlings first become familiar with possible underground hibernacula during the summer, and then during late summer or fall, bats might choose hibernacula based on social cues such as swarming conspecifics at the entrance of underground sites. In *Myotis* bat species, swarming can facilitate information transfer, mating, or both^2,15–17^. In Central-European *Myotis myotis*, swarming at potential hibernacula peaks between mid-August and mid-September (two to six weeks after we tracked bats at hibernacula), with many yearlings present^9,18^ suggesting that swarming contributes to the process of acquiring information about hibernacula by young bats. This hypothesis is further supported by the observation that mating has rarely been observed during swarming in *Myotis myotis*, suggesting that the primary function of swarming does not seem to be mating in this species^19^. Instead, recent evidence suggest that males exhibit a lek-mating system^8^, aggregating and remaining at mating roosts that might be located inside buildings, tree holes or bat boxes but also at underground sites, which are used and defended across years and visited by females^8,20^.

### Maternity colonies are social information centers

Our findings contribute to resolving a long-standing controversy in the life history of temperate bats: bats acquire information about hibernacula at their summer roost and remember those sites months later. Maternity colonies therefore have importance to bat populations beyond their function in reproduction: they are “information centers”^21^ that facilitate the transmission and updating of socially acquired knowledge about critical resources such as hibernacula. *M. myotis* is a long-lived animal with a reported maximum age of 37 years^9,22,23^. Such a long lifespan provides ongoing opportunities to gather information about new roosting sites and update knowledge about familiar locations, and summer maternity colonies provide adult females with ample opportunities to learn by following others to remote and previously unknown sites^24^. For *M. myotis*, updating knowledge on suitable hibernacula can be lifesaving because any given winter site can become unusable or inaccessible for that winter. As climate change and other anthropogenic influences alter resources used by bats^25,26^ access to social information may be an important mechanism for responding to rapid environmental change^27^.

Social learning about bat hibernacula is a hypothesis that has been notoriously difficult to test until now, as it requires a massive sampling effort given the rarity of observing such co-movements, the large number of hibernation sites, and the distances between them. Our evidence of socially transmitted knowledge of hibernation sites in bats, based on analyses of 38 years of monitoring efforts (frequently involving citizen volunteers), highlights the unique value of long-term datasets, which can provide rare insights into behavioral processes that are otherwise challenging to detect, particularly in cryptic species such as bats. Such datasets not only provide insight into the social structure of animal societies and the adaptive benefits of group-living, they are also critical for developing effective conservation strategies in a rapidly changing world^28,29^.

## Lead contact

Simon Ripperger

## Materials availability

This study did not generate new unique reagents.

## Acknowledgements

We thank Isabelle Waurick and Frieder Mayer for support during lab work. This work would not have been possible without the help of a large number of bat-enthusiast volunteers who dedicated countless days and hours of their free time to generate this unprecedented dataset. We are particularly grateful to Dr. J. Haensel, who was a pioneer of bat banding in Germany and without whom this monitoring program would likely not exist in its current form today. In addition, we thank P. Benda, J. Berg, J. Blesznowska, T. Blohm, K. Bogon, B. Borowski, Colelctive F. Bongers, N. Brunkow, O. Buexler, P. Busse, J. Cichocki, U. Damm, J. Dekker, C. Dietz, U. Dingeldey, Collective D. Dolch, K. Drabinska, Fam. Dronski, M. Dziegielewska, F. Ehlert, Mr. Fiedler, T. Frank, K. H. Froede, J. Froemert, J. Furmankiewic, K. Genz, H. Gille, M. Globig, A. Glover, M. Goettsche, N. Goldmann, J. Gorniak, Collective A. Griesau, E. Grimmberger, A. Hagenguth, ASG Hahneberg e.V., M. Hammer, Mrs. Handschak, D. Hargreaves, M. Heddergott, U. Heise, U. Hermanns, U. Hoffmeister, D. Horacek, J. Horn, A. Iben, M. Ignaczak, J. & J. Jablonski, C. Kallasch, Dr. M. Kares, D. Karoske, A. Kepel, Fam. Kiesewetter, J. Klawitter, C. Kuthe, R. Koch, T. Kokurewicz, Mr. Kopp, C. Kronmarck, Dr. R. Labes, Mr. Lappin, M. Lehnert, T. Liebscher, O. Lindecke, D. Linton, U. Loeser, G. Maetz, E. Malle, Mrs. Manzke, H. Maternowski, H. Matthes, F. Meisel, F. Meyer, H. Miethe, N. Moritz, R. Moritz, G. Nessing, oeKO-LOG Freilandforschung, Office K & S Berlin, B. Ohlendorf, W. Oldenburg, G. Pelz, H. Pommeranz, S. Petrick, A. Petzold, G. Preschel, W. Rackow, S. Rosenau, M. Rossner, Collective Rostock, W. Sauerbier, K. Sachanowicz, O. Schaefer, A. Schewe, W. Schober, Dr. A. Schmidt, C. Schmidt, H. Schroeder, H. Schuett, J. Schulenburg, M. Schulte, N. Starik, T. Staudt, D. Steinhauser, S. Stephan, C. Stichel, T. Teige, C. Telatynski, R. Tismer, J. & J. Teubner, C. Teumer, J. Thiele, S. Tost, S. Trost, Z. Urbanczyk, F. Verhaeghe, A. Vollmer, S. Voss, G. Walczak, M. Wilhelm, W. Willems, A. Woiton, G. Wojtaszyn, A. Zapart, J. Zejmo, and U. Zoephel

## Author contributions

Conceptualization, S. R., G. C., and A. A. F.; formal analysis: S. R. and C. G. investigation: S. R., A. A. F., L. I., J. H., B. K., R. H., K. D., and P. V.; writing – original draft, S. R., G. C., and A. A. F., Funding acquisition: S. R.

## Supplementary results

### Sampled bats

We analyzed 30,882 observations of the presence of 13,852 individually marked bats (7369 females, 6377 males, and 106 bats with unknown sex) recorded from January 6, 1985 to February 17, 2023 (38 years). Of these marked bats, 6412 were seen more than once, and 184 bats were seen at least 10 times (Fig. S1).

**Figure S1.**
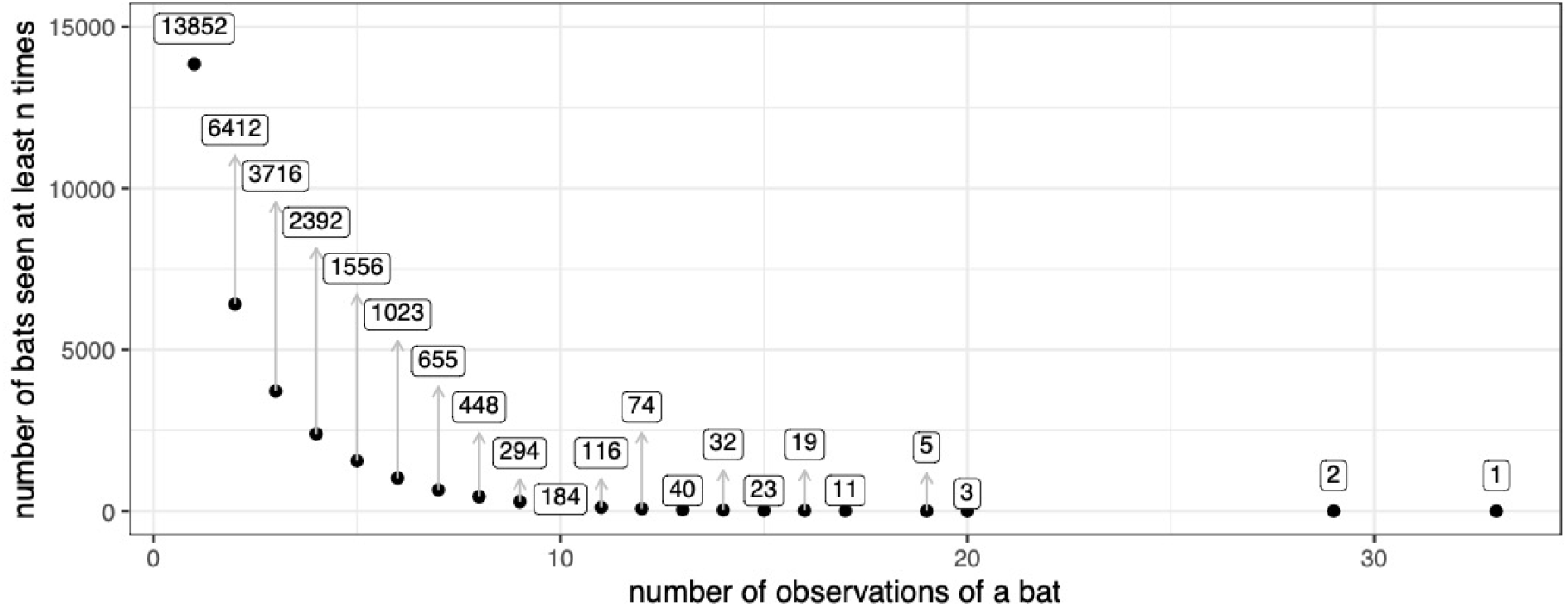
Number of repeated observations per bat. Points show the number of bats (Y-axis) that were seen at least n times (x-axis). Labels show the exact count of bats on the y-axis.

### Female bats used up to 4 and males up to 5 sampled hibernation sites

We found that 3421 females were observed at 1-4 winter sites (mean=1.18, 95% highest density interval (HDI): 1.16 to 1.19) and 3427 males were observed at 1-5 winter sites (mean=1.16, 95% HDI=1.14 to 1.17). After accounting for sampling effort, the average difference in counts between sexes (females - male) is only 0.03, 95% HDI=0.01 to 0.05 (Fig. S2). This difference was caused by females being more likely to switch hibernation sites (Fig. S4).

**Figure S2.**
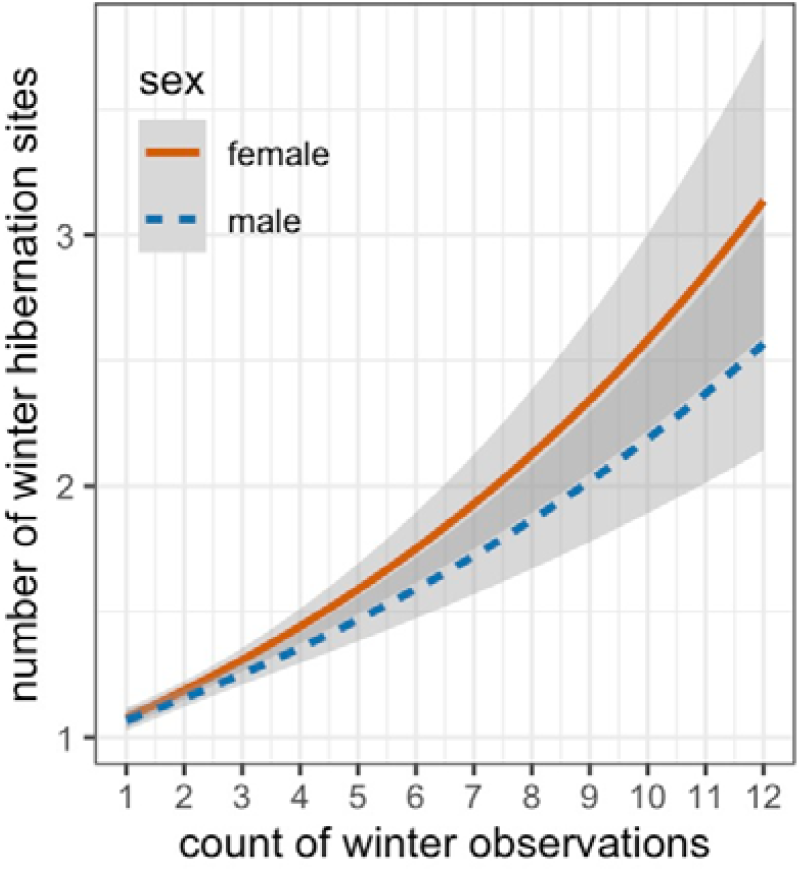
Not all winter hibernation sites were sampled. Lines show expected number of winter hibernation sites plotted by sampling effort (x-axis) for females (orange line) and males (blue dashed line). Females were seen at more of the sampled hibernation sites per unit of sampling effort.

### Sampled roosts

Bats typically move from a summer to a winter site. We sampled 351 roost sites that were used as winter hibernaculum (n=227), summer maternity (breeding) colonies (n=103), summer non-maternity colonies (n=122), and fall sites used for mating (“swarming” sites, n=15). Most sites (285) were used only during the summer or winter, but 65 sites were used during summer and winter seasons. For cases with known sex, winter observations of bats at hibernaculum (42% of observations) were 48% female (6166 female, 6596 male), summer observations at maternity (breeding) colonies (50% of observations) were 76% female (11473 female, 3621 male), and summer observations of non-maternity colonies (7%) were 65% female (1456 female, 777 male, Fig. S3). There were also 148 observations (0.5%) of bats at swarming sites, which were 45% female (66 female and 82 males). Bats were observed at up to 5 winter hibernation sites but gaps in their location show that they also used some unknown number of unobserved winter hibernation sites. There were 38,299 opportunities to see a marked bat in the winter (a possible winter observation), and 33% (12796 cases) of these possible winter observations were at known (sampled) sites. The average probability of a marked bat being observed in a winter was 36.8% (n=13,582). If a bat was seen at least once at a known hibernation site (n=6,875), the average probability of it being observed in a given winter was 74%.

**Figure S3.**
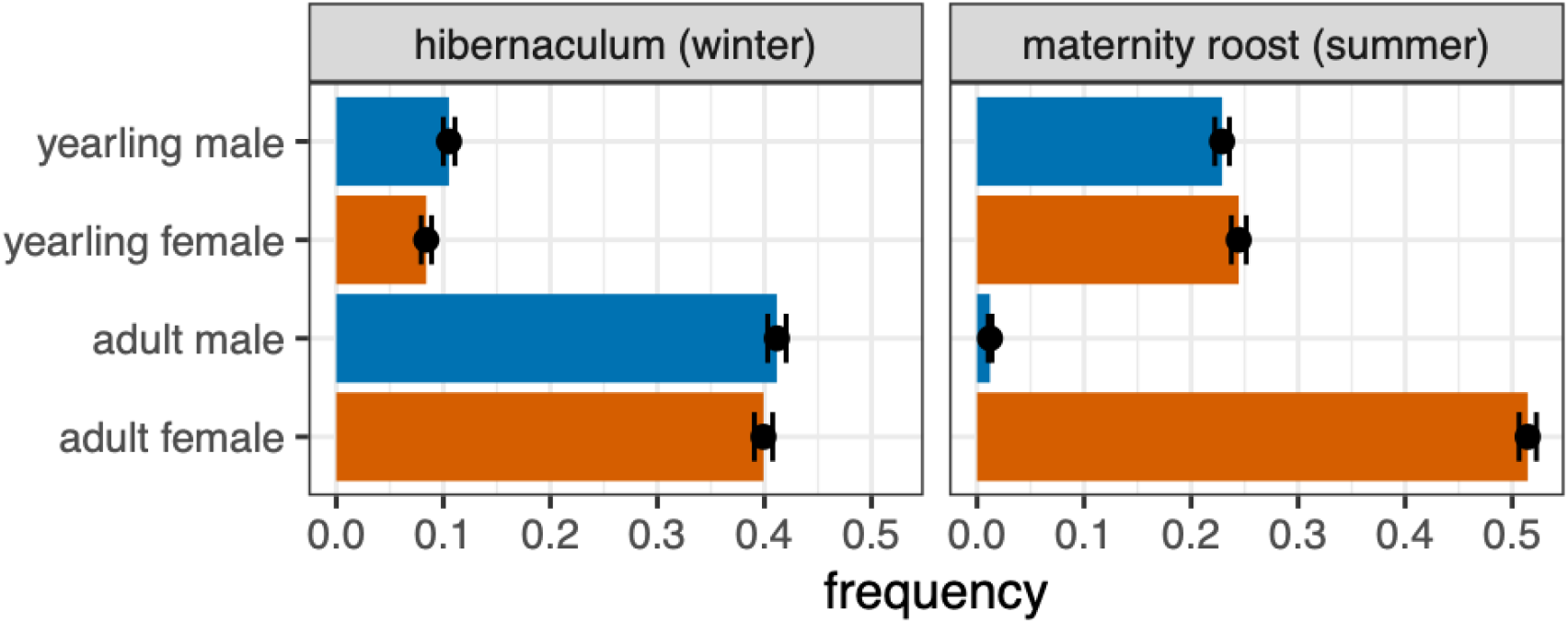
Sex and age composition by roost type. Error bars are exact 95% confidence intervals from binomial test.

**Figure S4.**
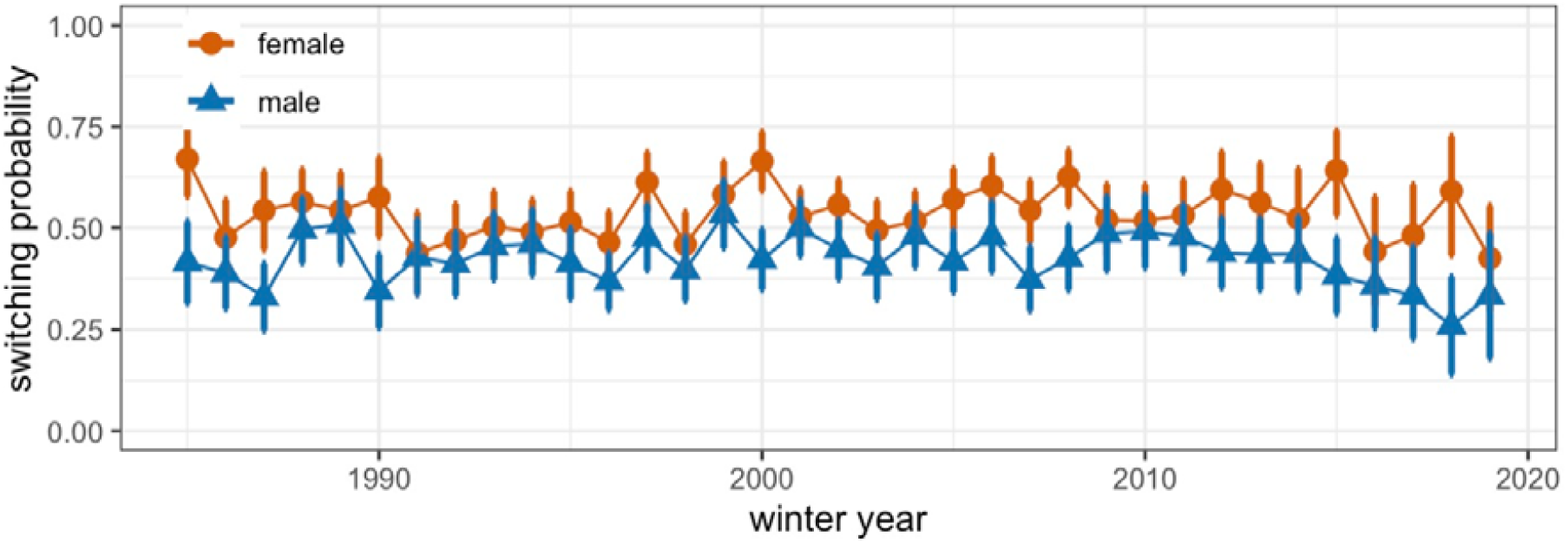
Mean probability of switching hibernation sites over time for females (orange circles) and male bats (blue triangles). Error bars are bootstrapped 95% confidence intervals.

### Bats that share summer roosts are more likely to share winter hibernacula

During the summer, the average number of possible co-roosting associations across pairs was 5.9 (range 1-16), 79% of pairs were never seen together, and the average co-roosting rate was 8.6%. During the winter, the average number of possible associations across pairs was 4.9 (range 1-16), 91% of pairs were never seen together, and the average co-roosting rate was 2.8%.

**Figure S5.**
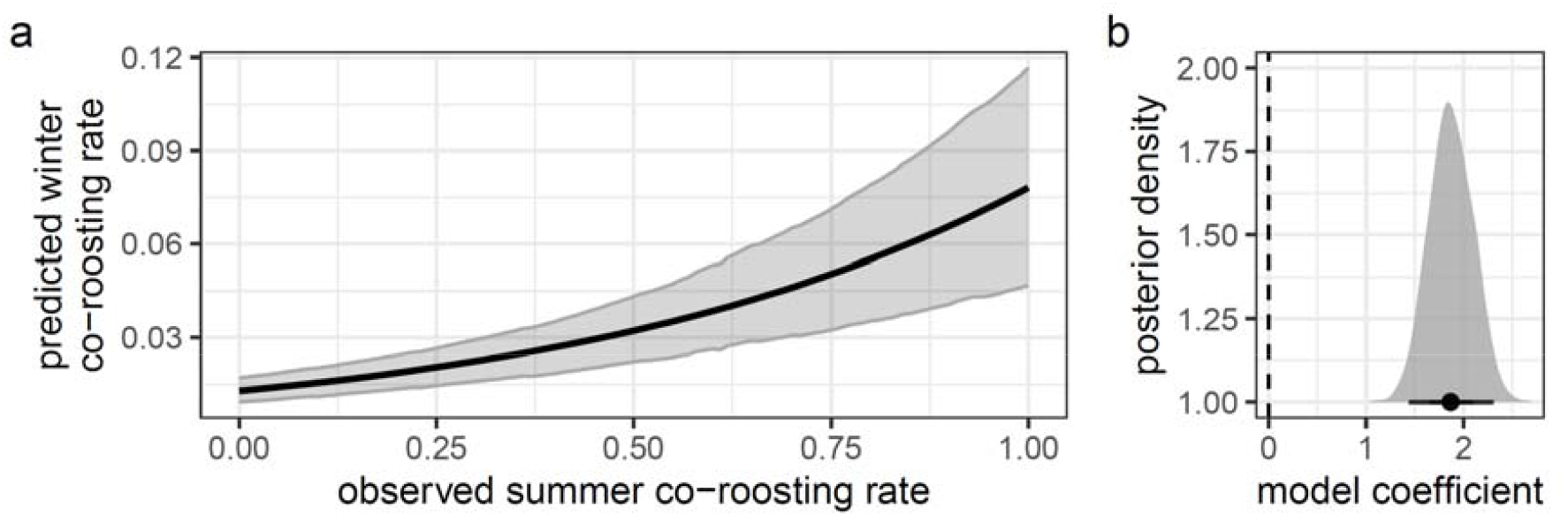
Observed summer co-roosting rates predict observed winter co-roosting rates. Panel A shows predicted probability of winter associations based on observed summer co-roosting rates. Shaded region is 95% credible interval. Panel B shows the posterior distribution of the model coefficient for summer co-roosting rate.

**Figure S6.**
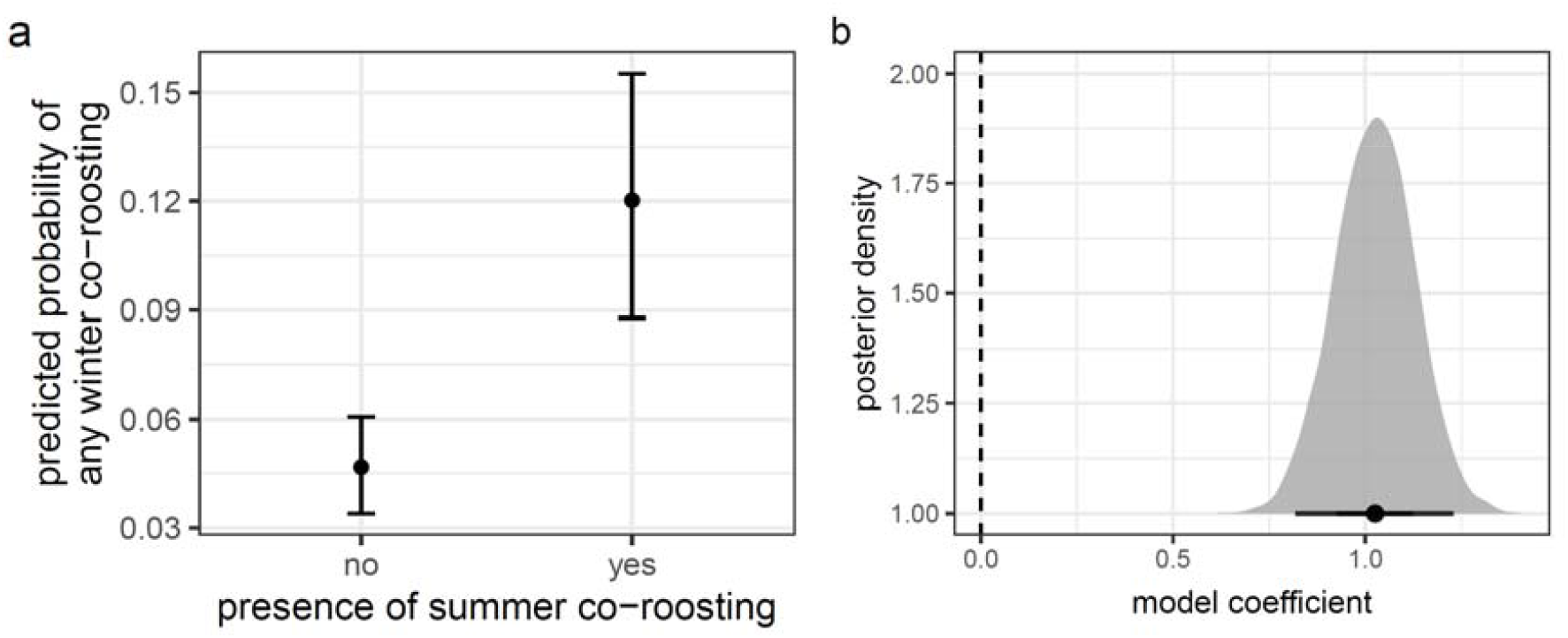
Bat pairs never seen together during the summer are less likely to be observed together during any winter. Panel A shows predicted probability of winter co-roosting based on presence or absence of summer co-roosting rates. Error bars are 95% credible intervals. Panel B shows the posterior distribution of the model coefficient for summer co-roosting rate.

**Figure S7.**
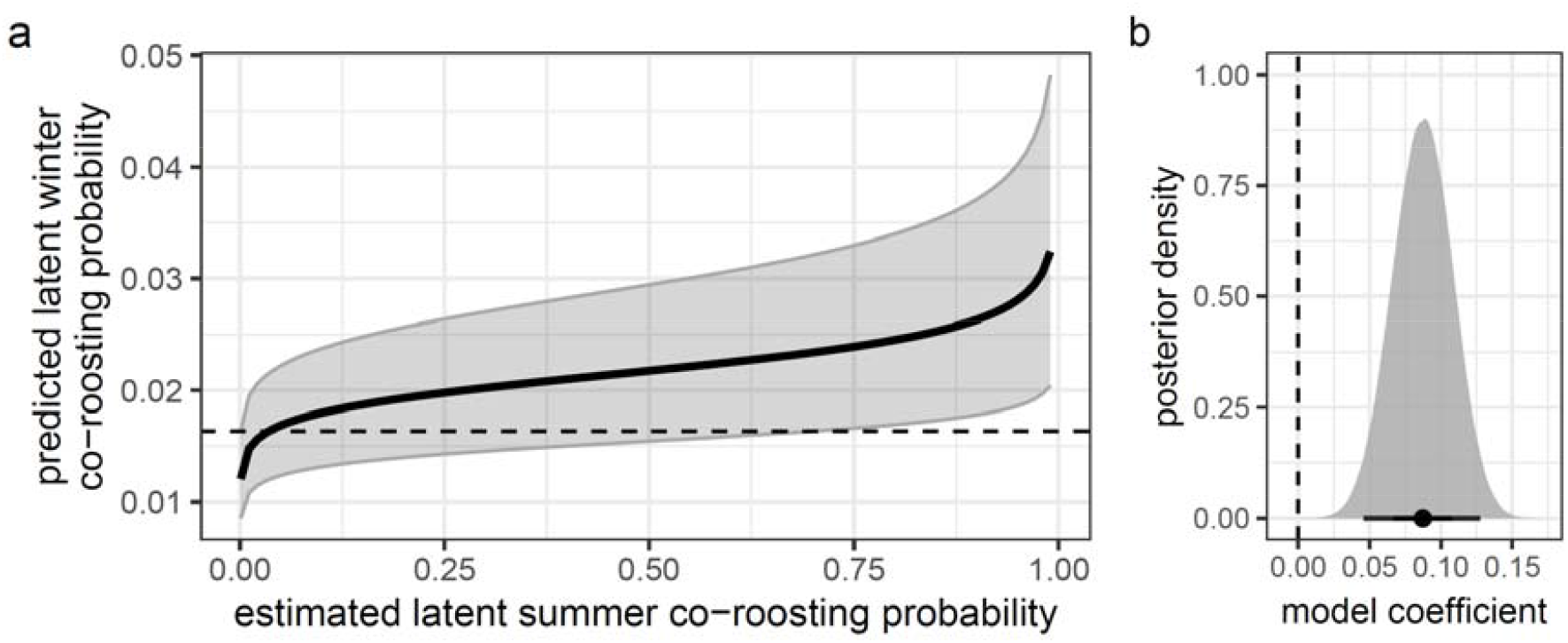
Latent winter and summer co-roosting rates are correlated. Panel A shows estimated probabilities of winter co-roosting by estimated probability of summer co-roosting using the bison R package. Shaded region is 95% credible interval. Dashed line is upper 95% credible interval for summer co-roosting probability of 0.001. Panel B shows the posterior distribution of the model coefficient for summer co-roosting rate.

**Figure S8.**
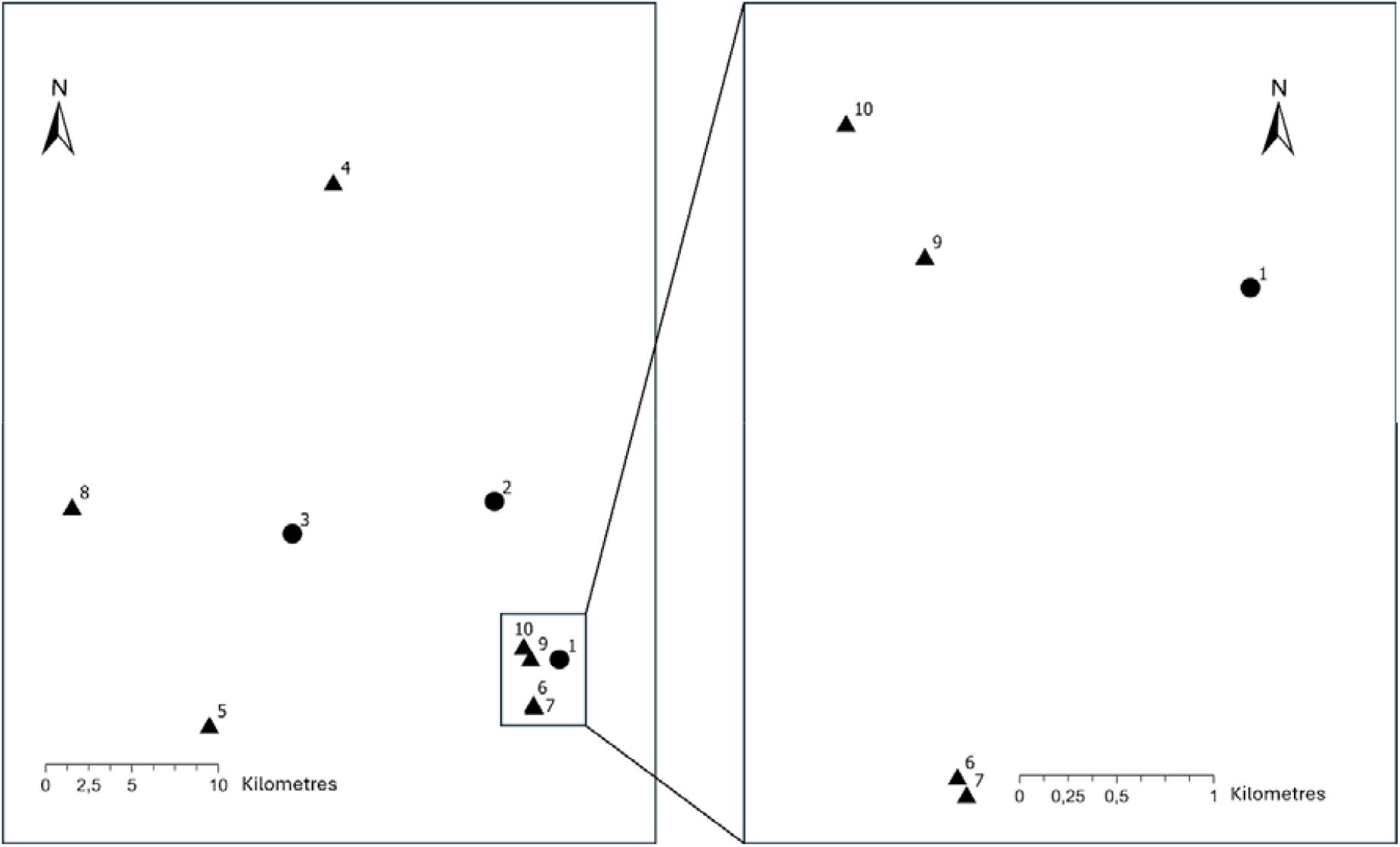
Map of associations detected by automated telemetry. Relative locations of three maternity colonies (circles 1-3) and seven hibernacula (triangles 4-10) that were equipped with tracking ground nodes during July and August 2022. Bats shown in Figure 3 were tagged at the maternity colony 3 (left panel).

**Table S1.** Model coefficients for estimates of effect of summer co-roosting on winter association.

| Model | Term | Estimate | Lower 95% CrI | Lower 95% CrI |
| --- | --- | --- | --- | --- |
| Model 1 | intercept | -4.34 | -4.65 | -4.04 |
|  | **observed summer co-roosting rate** | **1.87** | **1.44** | **2.31** |
|  | multi-membership pair | 1.87 | 1.61 | 2.18 |
| Model 2 | intercept | -3.53 | -4.04 | -3.03 |
|  | **observed summer co-roosting (0/1)** | **1.02** | **0.82** | **1.23** |
|  | natural log of summer opportunities | -1.53 | -2.20 | -0.86 |
|  | natural log of winter opportunities | 2.02 | 1.46 | 2.61 |
|  | multi-membership pair | 1.78 | 1.52 | 2.08 |
| Model 3 | intercept | -3.76 | -4.10 | -3.42 |
|  | **estimated summer association** | **0.12** | **0.07** | **0.18** |
|  | multi-membership pair | 1.81 | 1.55 | 2.10 |

