## Supplementary material for "Socially transmitted knowledge of hibernation sites in bats": Methods

**EXPERIMENTAL MODEL AND STUDY PARTICIPANT DETAILS**

**Study species**

The Greater mouse-eared bat (*Myotis myotis*) is distributed across western, central, and southern Europe, with the south-eastern border of its range in Asia Minor^1^. The Greater mouse-eared bat is one of the largest species of *Myotis* (wingspan 350–430 mm). In Germany, maternity colonies usually form, often in buildings and attics, in late March or early April with highest numbers of adult females in mid-May. Births usually start in June and end at the beginning of July with considerable variation across years, colonies, and weather conditions. In late summer and fall, activity increases at underground sites such as caves, cellars, and mines. From August onwards, swarming can be observed at caves reaching a maximum mid-August to mid-September, and juveniles are present in large numbers. First torpid bats can be found as early as in October and hibernation may last up to five months, leaving juveniles only about five months for their development before winter. Hibernating *M. myotis* often roost alone, but mixed-sex groups include up to several hundred bats have occasionally been observed (reviewed in Zahn et al. (2022)^2^).

**Research permits**

Necessary research permits were obtained by Landesamt für Umwelt, Brandenburg, and Landesamt für Arbeitsschutz, Verbraucherschutz und Gesundheit LFU-N4-4730/22+55#238583/2022, LFU-N4-4730/22+23#166282/2022, 105-N4-4730/24+49#255433/2023, and LAVG-V2-2347/77+1#8365/2022).

**METHOD DETAILS**

**Summer and winter census**

All banding data are deposited center for bat banding at the Saxon State Office for Environment, Agriculture and Geology in Dresden, Germany, which is one of the two national centers for bat banding and that curates a bat banding database and distributes bat bands for long-term monitoring projects. Most data were collected in the course of a long-term monitoring in the states of Brandenburg and Berlin, Germany, which have a long tradition of bat banding. Dr. Joachim Haensel and associated volunteers contributed most data that were the basis for this study until his death in 2014. Their work was continued by the volunteer groups “Mausohr e.V.” and “Berliner Artenschutz Team – BAT – e.V.” and other volunteers that coordinate annual census.

For banding *Myotis myotis*, aluminum bat band type A were used. In addition to the letter that indicates the type, each band carries an individual number. The weight of the band is less than 0.1 g.

We analyzed bat census data that comprised 30,882 observations of 13,852 bats at summer, fall, or winter roosts from January 1, 1985 to February 17, 2023. Most summer roosts are inside buildings or overground technical structures (e.g., bridges), and most hibernacula are in underground structures such as cellars, bunkers etc. The median number of hibernacula that were visited per year was 34 (max = 56 hibernacula). During the winters of the SARS-Covid 2 pandemic, only 3-6 hibernacula were counted. The median number of maternity colonies counted per year was 9 (range: 2-20). During observations, band numbers were recorded and bats were sexed. Unbanded bats were banded during both summer and winter.

**Tracking of appearances at hibernacula**

In 2022, we fitted 40 individuals of *Myotis myotis* from three maternity colonies with Dulog sensor nodes (see Morosse et al. 2025^3^ for technical details) that use the functionality of the BATS wireless biologging network^4^. Animal-borne nodes were housed inside fingertips of small-sized nitrile lab gloves and glued between the scapulae of each bat using Sauer latex skin adhesive. Total weight of housed nodes was below one gram. The bat with a lowest body mass (one juvenile) weighted 19.5 g, all other bats were 20 g or more in mass. Consequently, tags did not exceed 5 % of the bats body weight.

Usually, these animal-borne nodes serve as proximity sensors that log encounters between tagged individuals inside (e.g. Ripperger et al. 2019^5^) and outside the roost (e.g. Ripperger et al. 2021^6^), and data are downloaded remotely to base stations that are positioned at roosts. However, the download option malfunctioned, so we analyzed contacts to base stations at seven hibernacula. These hibernacula were regularly used for hibernation by bats from the three maternity colonies bats where they were banded.

Bats were tagged on July 19^th^ of 2022 at three different maternity colonies (Sites1-3 in Figure S8). Adult females and yearlings were collected by hand from their roosting sites inside the maternity colonies. To target mother-offspring pairs, we selected and tagged adult bats and pups that roosted in close proximity or that were attached to each other. Genetic analyses (described below) confirmed the suspected mother-pup relationship.

Nodes broadcasted beacon signals every eight seconds between 9 pm and 6 am of each night and contained an individual ID that allows one to attribute a signal to an individual bat. Beacon signals were stored by the base station along with the ID, the RSSI (received signal strength indicator), and a time stamp obtained from a GPS module. Base stations were positioned inside hibernation sites to minimize risk of theft. Since most hibernacula were bunkers made from massive ferro concrete base stations could not update initial time stamps due to lacking reception by GPS satellite signals. Therefore, absolute time stamps might be off by several minutes to hours after a runtime of several days, however relative timestamps that served to infer time differences between arriving bats at hibernacula remain credible. Inter-beacon intervals of individual bats that were documented at base stations usually equaled eight seconds. The last beacon signals broadcasted by bats were received on August 4^th^ 2022 (maximum recorded runtime of 16 days).

**Genetic analyzes to assign mother-pup pairs**

Tracking data suggested that two juveniles conducted tandem flights with one adult female. To check whether the adult bat was the mother of either or of both juveniles (birth of twins are rare in *Myotis myotis* but have been observed (Zahn et al. 2022)), we genotyped 20 microsatellite loci. Tissue samples were obtained from the wing membrane of all tagged bats using 4 mm wing biopsy punch. Samples were stored in 80% ethanol. Genomic DNA was extracted using a salt–chloroform procedure. Fragments were amplified using uniplex PCR with M13 tails^7^ (fluorescent labels FAM, Yakima Yellow, Atto 550 (NED), Atto 363 (PET)). We used the SeqStudio™ Genetic Analyzer with SeqStudio™ Genetic Analyzer Cartridge v2 (both Applied Biosystems™/ ThermoFisher Scientific) for genetic analyses. For sizing, we used the GeneScan 500 LIZ Size Standard (Applied Biosystems™/ ThermoFisher Scientific).

**Software**

We used QGIS Desktop 3.44.8 (Figure 1) and ArcGIS Pro 3.3.2 © 2024 Esri Inc. (Figure S8) to create maps. Statistical analyzes were run in R (version 4.4.3; R Core Team, 2025^8^).

**QUANTIFICATION AND STATISTICAL ANALYSIS**

**Sex bias in the use of hibernacula**

To predict the number of winter hibernation sites used by bats of each sex, we fit a Bayesian hurdle Poisson model using the brms R package^9^ and extracted predictions using the marginaleffects R package^10^. Along with sex, we included the log number of winter observations as an offset term to control for sampling effort per bat. We assessed model fit with posterior predictive checks.

**Switching of hibernacula across winter**

To model how often bats switched hibernation sites between winters, we first compiled 9045 cases of possible switches where a bat was seen at a known winter site and was either seen there the last winter (i.e. it returned to the same winter site) or not (i.e. it switched winter sites). We then fit a Bayesian multilevel Bernoulli model with population-level (fixed) effect of sex and group-level effects (random or varying effects) of bat and winter season using the brms R package. We estimated predictions using the marginaleffects package. We assessed model fit with posterior predictive checks.

**Distance between birth site and first and second hibernaculum**

We found 52 bats (20 males and 32 males) with known birth sites (i.e. they were seen as yearlings at a summer roost) that used two different hibernation sites in their first and second winter. We then estimated the probability that their first hibernation site was closer to their birth site compared to the next hibernation site where we observed them. To do this, we fit a Bayesian Bernoulli model in brms with sex as a predictor of the outcome that the first hibernaculum sighting was closer than the second. To test the null hypothesis that distances to hibernation site were not closer in the first year, we also used a nonparametric permutation test where we randomized which years each bat was seen at the hibernation sites they used. This null model accounts for the lifetime home range of the bat and the nonrandom distances of the sampled sites.

**Co-roosting of yearlings and adults at hibernation sites**

In 931 observed movements of yearlings from the maternity colony to the first hibernaculum, 364 occurred with an adult roostmate. We compared this observed proportion of cases (0.39) to values expected from a null model simulating co-movements expected by chance (without social information transfer). For this null model, we randomized the dates of appearance within each individual bat and type of site (e.g. winter hibernaculum, summer maternity colony, other summer roost). For example, if a bat was seen at hibernation site A in year 1, A in year 2, B in year 3—then one permuted sequence could be that it used site B in year 1 and site A in years 2 and 3. In this null model, bat movements are therefore independent of each other.

**Co-roosting probabilities in summer and winter**

We counted actual and possible co-roosting associations in summer roosts and in winter hibernacula for the 180 bats that were seen at least 3 times in the winter and at least 3 times in the summer. To get possible associations, we created a possible observation period for each bat as the period between our first and last observation of it, and we assumed that two bats could have been seen together for any observation date where both bats could have been observed.

For the first beta-binomial model, the outcome was the count of winter co-roosting observations out of the total number of winter opportunities to co-roost, the predictor was the observed summer co-roosting rate (the proportion of summer opportunities in which the dyad was observed together), and a multi-membership group effect accounted for the non-independence of dyads sharing one or both bats. We used default uninformative priors. Rhat values for model parameters were 1.00-1.01 and effective sample sizes were 837 to 4061.

For the second model, we asked whether pairs seen together during any sampled summer (true/false) were more likely to be seen together during any sampled winter (Bernoulli outcome). To control for sampling effort, we included log counts of possible summer co-roosting and log counts of possible winter co-roosting for each dyad as covariates. We used default uninformative priors. Rhat values for model parameters were 1.00 and effective sample sizes were 3795 to 21,145.

For the third model, we built a joint model to account for uncertainty in estimated summer co-roosting rates using the Bayesian Inference of Social Networks (BISoN) framework in the bisonR package^11^. This approach propagates uncertainty in summer co-roosting rates into a regression predicting winter co-roosting rates. In the first two models, summer associations of 0/1 and 0/10 are treated as equivalent zeros, but in this third model, we replaced these point estimates with probability distributions by first using the actual and possible observations to fit a Bayesian social network edge-weight model. Rather than estimating a single co-roosting rate value for each dyad, this model estimated a posterior distribution describing the plausible range of each dyad's latent summer co-roosting probability. However, this model relies on imputation based on additional assumptions about the latent summer co-roosting rates underlying the observed counts. For priors on edge weights, we used a gaussian prior centered at -3 on the logit scale (corresponding to an co-roosting probability of 4.7%) with a standard deviation of 3, indicating that we expect most dyads to have low co-roosting probabilities, but with high variation. After estimating posterior distributions on summer co-roosting rates, we then passed these estimates into a beta-binomial regression model in which the response was the number of actual winter associations out of possible cases. The BISoN framework propagated uncertainty in the estimated summer co-roosting rates into the regression by repeatedly sampling from the posterior distribution for each dyadic co-roosting rate and fitting the regression across multiple imputed datasets. Parameter estimates and credible intervals therefore incorporate uncertainty arising from both the estimation of summer co-roosting rates and the relationship between summer and winter co-roosting rates.

**ADDITIONAL RESOURCES**
